# Software Toolkit for Fast High-Resolution TMS Modeling

**DOI:** 10.1101/643346

**Authors:** Sergey N. Makarov, Gregory M. Noetscher, Edward H. Burnham, Dung Ngoc Pham, Aung Thu Htet, Lucia Navarro de Lara, Tommi Raij, Aapo Nummenmaa

## Abstract

Computational modeling of Transcranial Magnetic Stimulation (TMS) is a tradeoff between computational speed vs. spatial precision. In this study, we introduce a software toolkit for high-resolution TMS modeling, which may offer the best of both. The toolkit employs the recently developed boundary element fast multipole method (BEM-FMM) with accurate solution computations close to tissue interfaces. It operates within the MATLAB platform and is designed for a broad, medically-oriented computational research community. To enable easy, subject-specific operation, the package is compatible with human head models generated with automated tools such as SimNIBS and FreeSurfer.

Both coil design in free space as well as efficacy and focality of TMS for a specific subject could be modeled by the package, including optional user-defined parametric loops. Detailed and widely used coil models, generated in both CAD and wire formats, may include several hundred thousand elementary current elements and observation spaces with approximately 1 M field points; the corresponding computational times are on the order of 1 sec. Detailed head models with approximately 1 M triangular facets and a mesh resolution of 0.6 points per mm^2^ are simulated in approximately 1.5 min which is arguably the fastest time to date. Further reduction of computational times is foreseen. The toolkit is augmented with a population of 16 ready-to-use head models for performing simulations for computational studies that do not involve MRI data collection. The toolkit is also augmented with a coil geometry generator capable of creating accurate coil models.

## 1. Introduction

For all three chief neurostimulation modalities – transcranial magnetic stimulation (TMS), transcranial electric stimulation (TES), and intracortical microstimulation (ICMS) – simulating the electric fields within a patient-specific head model is the major and often only way to foster spatial targeting and obtain a quantitative measure of the required stimulation dose (Bikson et al., 2018). Specifically, the net TMS E-field consists of primary and secondary components. The primary component – the E-field directly induced by the coil – can be determined using scalar and vector potentials, which are calculated based on the geometric features of the TMS coil. The secondary component – due to induced charges on tissue interfaces – can either be found using the boundary element method (BEM), the finite element method (FEM) or the finite difference method (FDM).

The above three methods differ in terms of computational speed and spatial precision. The boundary element method is fast with low-resolution models but its speed quickly deteriorates towards higher-resolution models. Salinas et al., 2007 and Salinas et al., 2009 have written two fast computer programs in C++ and interfaced them with MATLAB. The first program (3D coil generator) simulates a coil’s geometry; the second program (E-field generator) performs numerical calculations of the induced E-field in the volume of interest. The Helsinki BEM MATLAB library (Stenroos et al., 2007) and its refinements (Stenroos & Sarvas, 2012; Stenroos & Nenonen, 2012; Stenroos, 2016; Stenroos & Nummenmaa, 2016; Koponen et al., 2017) have been used for the purpose of TMS modeling as well. Different mathematical BEM formulations can be employed for TMS modeling (Gomez et al., 2018a).

The chief finite element TMS solver is the well-known, open-source transcranial brain stimulation modeling software SimNIBS (Thielscher et al., 2015; Opitz et al., 2015; Nielsen et al.,2018), whose most recent version, v2.1 (Saturnino et al., 2018), currently uses the open-source FEM software getDP. There are also many general-purpose, open-source solvers for finite element modeling, from high-level environments such as getDP (Dular et al., 1988), Deal.II (Bangerth et al., 2007), and FEniCS (Logg et al., 2012), to lower-level environments such as PETSc (Balay et al., 2018). Those solvers provide a practical and well-tested choice for creating problem-specific software solutions.

Commercial FEM software has options for adaptive mesh refinement and for a variety of accurate CAD coil geometries but tends to be slow compared to SimNIBS. Several studies (Miranda et al., 2003; Deng et al., 2011; Cho et al., 2010; Makarov et al., 2016; Wang et al., 2018) have used the commercial FEM solver ANSYS Maxwell 3D. Silva et al., 2008 and Salvador et al.,2011 used the commercial FEM solver COMSOL. Deng et al., 2013 used the FEM package MagNet (Infolytica, Inc., Canada) for a comprehensive focality study of 50 different TMS coil designs.

Multigrid fast and efficient FDM TMS simulators have been developed by Laakso & Hirata,2012 and Laakso et al., 2018. The commercial FDM software SEMCAD X was used for TMS modeling by Rastogi et al., 2017.

The problem with BEM, FEM, and FDM, is that they offer compromises between spatial accuracy and computational speed. Here, we use a new TMS modeling method, based on the coupling the boundary element method and the fast multipole method (BEM-FMM; Makarov et al., 2018; Htet et al., 2019), that offers both speed and accuracy at the same time. This approach is an integration of the BEM formulation in terms of surface charge density and FMM invented by Rokhlin and Greengard (Rokhlin, 1985; Greengard & Rokhlin, 1987; Gimbutas & Greengard,2015). The algorithm still possesses the major advantage of the conventional BEM – high speed – but is simultaneously capable of processing a very large number of surface-based unknowns. The BEM-FMM approach has demonstrated faster computational speed (10-1000 times faster) and superior accuracy for high-resolution head models as compared to both the standard boundary element method and the finite element method used in SimNIBS or commercial FEM software ANSYS Maxwell 3D (Htet et al., 2019). These advantages of the BEM-FMM approach have been recently independently and thoroughly confirmed (Gomez et al., 2018).

The present study introduces the open-source software package for high-resolution modeling TMS fields via BEM-FMM. The computational platform is basic MATLAB^®^ starting with version 2018a and with major FMM subroutines and other critical numerical subroutines converted to MATLAB executables, for both Windows and Linux. The software package is augmented with a population of sixteen high-resolution 2 manifold CAD compatible head models (Htet et al., 2018) with the edge length of cortical surface meshes of 1.4 mm from the Human Connectome Project (Van Essen et al., 2012). It is also augmented with a flexible coil geometry generator and a base collection of ten different coil design examples. Appendix A contains a concise user’s manual.

## 2. Materials and Methods

### 2.1 Numerical algorithm

#### Fast multipole method (FMM)

We integrate and use the open-source FMM library FMMLIB3 (FMMLIB3, 2017; Gimbutas & Greengard, 2015). In this version, there is no *a priori* limit on the number of levels of the FMM tree, although after about thirty levels, there may be floating point issues (L. Greengard, private communication). The required number of levels is determined by the maximum permissible least-squares error or method tolerance, which is specified by the user. The FMM is a FORTAN program compiled for MATLAB. The tolerance level `iprec` of the FMM algorithm is set at 0 or 1 (the relative least-squares error is guaranteed not to exceed 0.5% or 0.05%, respectively). This FMM version allows for a straightforward inclusion of a controlled number of analytical neighbor integrals on triangular patches to be precisely evaluated as specified below.

#### Boundary element method

We use the surface-charge approach mathematically called the adjoint double layer formulation (Rahmouni et al., 2018). Here, the unknown quantity is the charge density *ρ*(***r***′) induced at conductivity boundaries. The corresponding integral equation is obtained by writing the total electric field ***E***^*t*^ in a form that considers the primary solenoidal field ***E***^*p*^ and a conservative contribution of the secondary induced surface charge density ***E***^*s*^, i.e.

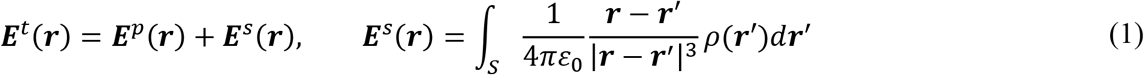

where *ε*_0_ is the permittivity of vacuum. Taking the limit of Eq. (1) as ***r*** approaches surface S from both sides (inside and outside) and using the continuity condition for the normal current component, *σ**E**^t^*(***r***), one obtains the adjoint double-layer equation (Barnard et al., 1967; Makarov et al., 2016; Rahmouni et al., 2018) in the form

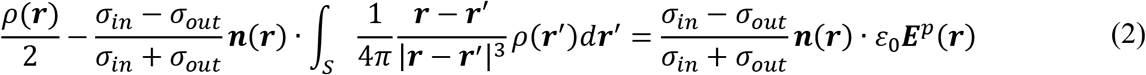

where ***n***(***r***) is the outer normal vector to the particular boundary and *σ_in/out_* are the two distinct tissue conductivity values on either side of the boundary. The first (simpler) task of the numerical solution is to compute the excitation – the primary electric field ***E***^*p*^(***r***) – everywhere at tissue conductivity boundaries in Eq. (2). The second (more complicated) task is to compute the induced charge density by solving integral equation (2) of the second kind iteratively and thus compute the secondary electric field ***E***^*s*^. Both fields add up.

#### Computation of primary electric and magnetic fields

Primary electric and magnetic fields ***E***^*p*^ and ***B***^*p*^ of a TMS coil are computed using FMM. The number of elementary current filaments is ultimately unlimited. Consider a small straight element of current *i_j_*(*t*) with segment vector ***s***_*j*_ and center ***p***_*j*_. Its magnetic vector potential, ***A***^*p*^, at an observation point ***c***_*i*_ is given by (Balanis, 2012)

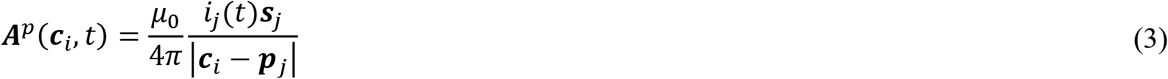

where *μ*_0_ is magnetic permeability of vacuum and index *p* again means the primary field. The electric field generated by this current element is

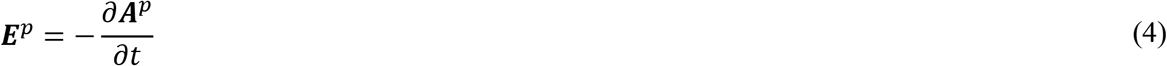

Assuming harmonic excitation with angular frequency ω and omitting the redundant phase factor of −*j*, one has

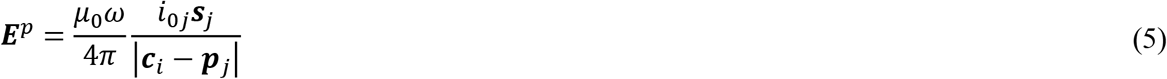

Now consider multiple straight short current segments with moments *i*_*j*0_***s***_*j*_ forming the coil. Also consider multiple observation points ***c***_*j*_ which coincide, for example, with triangle centroids of the head surface mesh. For every observation point, the electric field in Eq. (5) is computed via FMM as a potential of a single layer repeated three times, i.e., separately for each component of the field. The corresponding pseudo charges used in FMM function `lfmm3dpart` of the FMM library (FMMLIB3, 2017; Gimbutas & Greengard, 2015) are chosen in the form (to within a constant)

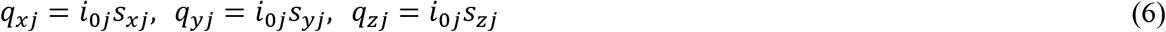

The same element of current generates a magnetic field ***B***^*p*^ given by Biot-Savart law

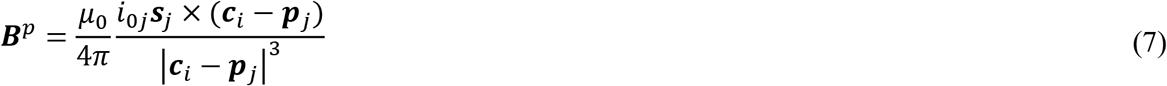

For multiple segments and multiple observation points, Eq. (7) cannot be evaluated using the FMM framework directly. However, it can be rewritten in the form

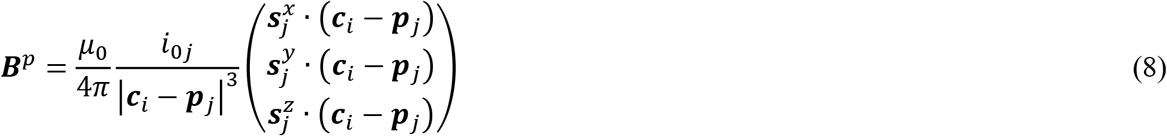

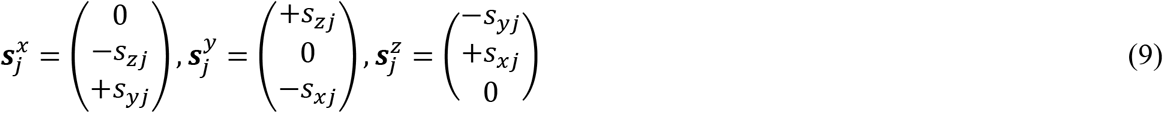

which is equivalent to the electric potential of a double layer (layer of electric dipoles). This potential is to be computed three times and with three different sets of pseudo dipole moments given by Eq. (9) and used in the FMM function `lfmm3dpart` (FMMLIB3, 2017; Gimbutas & Greengard, 2015). This is another standard FMM task.

#### Computation of secondary electric field

The fast multipole method (Rokhlin, 1985; Greengard & Rokhlin, 1987) speeds up computation of a matrix-vector product by many orders of magnitude. In the present problem, such a matrix-vector product appears when an electric field from many point sources *ρ*(***r***′) in space has to be computed at many observation points ***r***. Namely, discretization of the surface integral in Eq. (1) or Eq. (2) and use of the zeroth-order piecewise constant basis functions (pulse bases) yields

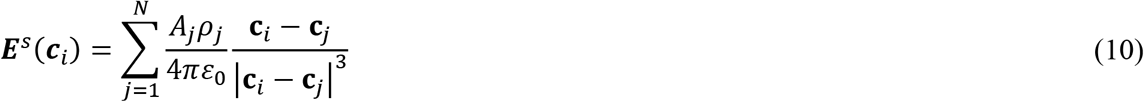

where *A_i_, **c**_i_, i* = 1,…,*N* are the areas and centers of triangular surface facets *t_i_; ρ_i_*, are triangle surface charge densities in C/m^2^, and *A_j_ρ_j_* are elementary charges located at **c**_*j*_. Eq. (10) is straightforwardly computed via the same FMM library, as an electric field of a given charge distribution. Such a computation is done at every step of an iterative solution for integral equation (2).

Approximation (10) is inaccurate for the neighbor facets. Using the Petrov-Galerkin method with the same pulse bases as testing functions, we employ a refined computation:

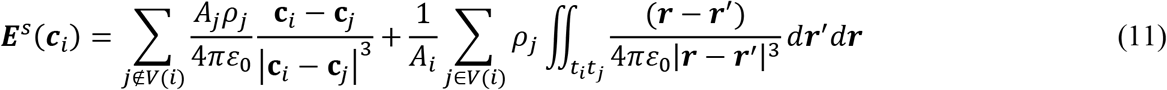

where *V*(*i*) encompasses a neighborhood of facets around observation triangle *t_i_*. The inner integrals in Eq. (11) are computed analytically (Wilton et al., 1984; Wang et al., 2003; Makarov et al., 2016) while the outer integrals use a Gaussian quadrature of 10^th^ degree of accuracy (Cools,2003).*V*(*i*) includes *K ≪ N* nearest triangular facets. Inclusion of a small number of precomputed potential neighbor integrals (three to sixteen) improves the convergence of the iterative solution; inclusion of a larger number has a negligible effect. The present BEM-FMM approach performs precise analytical integration over *K* = 12 – 16 closest neighbor facets.

The same approach is used for volume field computations, when an observation point is located close to the charged boundary. In this case, the contribution of charges on neighbor facets is accurately averaged as prescribed by the second term on the right-hand side of Eq. (11).

#### Iterative solution

Eq. (2) is solved iteratively using the generalized minimum residual method or GMRES (Barrett et al., 1994; Saad, 2003) implemented by Drs. P. Quillen and Z. Hoffnung of MathWorks, Inc. Its overall performance and convergence are excellent, especially for complicated head geometries.

#### Algorithm accuracy and speed

The accuracy of the algorithm has been tested using side-by-side comparisons with analytical solutions, the open-source FEM solver getDP employed by SimNIBS, and the commercial FEM software ANSYS Maxwell with adaptive mesh refinement (Makarov et al., 2018; Htet et al., 2019). Both spherical and anatomically realistic head models were tested. For high-resolution models, the BEM-FMM algorithm ran significantly (10-1000 times) faster than the FEM and provided a better solution accuracy(Htet et al., 2019).

For the head models described below with approximately 0.9 M facets and with an average edge length of 1.4 mm in the cortex, run times necessary to achieve a relative residual of 10^−3^ and obtain a good solution accuracy are about 90 sec (4.5 sec per iteration for 20 iterations or less). These data were compiled on an Intel Xeon E5-2698 v4 CPU (2.20 GHz) server, 256 Gb RAM, MATLAB 2018a. Lowering the relative residual does not improve accuracy.

### 2.2 Head models

#### Model format

Any head model comprised of *.stl (stereolithography) files for each individual brain compartment could be imported. Such models are routinely generated by SimNIBS (Saturnino et al., 2018) and FreeSurfer (Fischl, 2012). For example, any of the 50 head models from the Population Head Model Repository created by Lee et al., 2016 (see also Lee et al., 2018; The Population Head Model Repository, 2017) using SimNIBS software and the Connectome MRI Database may be used. In this case, the typical model has about 0.7 M triangular facets in total (skin resolution was lowered via post-processing).

#### Built in head models

The toolkit is augmented with a collection of sixteen high-resolution, 2-manifold CAD compatible head models within the MATLAB platform described in (Htet et al.,2018). Each model contains skin, skull, CSF (cerebrospinal fluid), GM (gray matter), cerebellum, WM (white matter), and ventricles head compartments in the form of triangulated manifold shells with nearly uniform triangle density. Every surface object can be 3D printed. The models are compatible with a variety of modeling software, including ANSYS Electronics Desktop, CST Studio Suite, Sim4Life/SEMCAD, and other electromagnetic simulation software packages.

The collection is based on MRI data for sixteen subjects from the Human Connectome Project (Van Essen et al., 2012) segmented using the SimNIBS processing pipeline (Saturnino et al.,2018). The MRI data were kindly provided by the Human Connectome Project WU-Minn Consortium (PIs: David Van Essen and Kamil Ugurbil; 1U54MH091657, see S1200 Reference Manual 2018). The average number of triangular surface facets in a model is 866,000, the average triangle quality is 0.25, the average edge length for all surfaces is 1.48 mm, and the average surface mesh density or resolution is 0.57 points per mm^2^. This mesh resolution (1 to 1.5 mm) corresponds approximately to the high-quality 3 T images.

#### Head model visualization and pre-processing

The head models are visualized in MATLAB either as a surface mesh for every brain compartment (Fig. 1a) or as head cross-sections (2-manifold polygons for skin, scalp, and CSF) in any principal plane – coronal (XZ), sagittal (YZ), and transverse (XY) – located anywhere in space. As an example, Fig. 1a shows white matter, gray matter, and CSF (inner skull) shells for Connectome subject # 110411. Next, Fig. 1b shows three cross-section cuts obtained for the same subject at *z* =-0.0185 *m, y* = 0, *x* = 0.01 *m*. Furthermore, the integral mesh characteristics, including mesh quality, can be computed.

**Fig. 1a.**
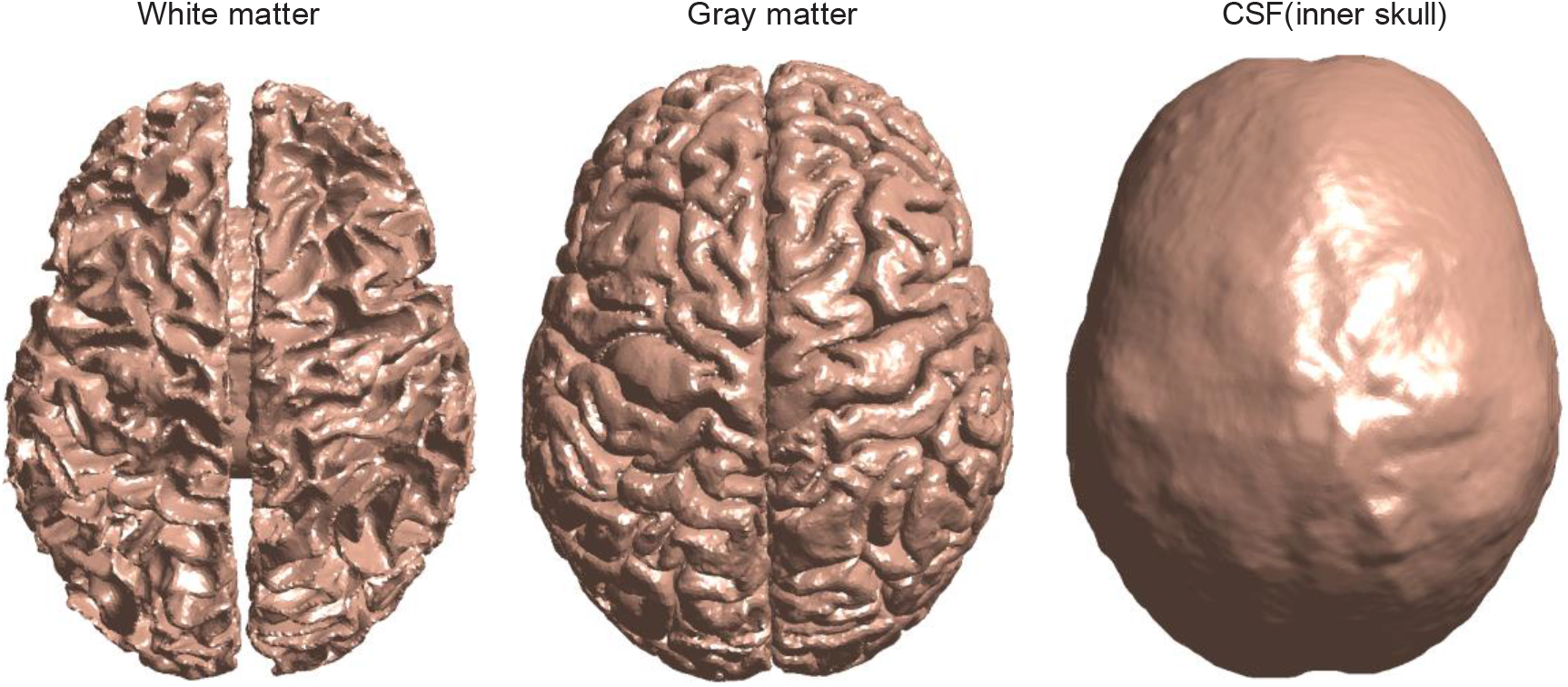
White matter, gray matter, and CSF (inner skull) shells for Connectome subject # 110411.

**Fig. 1b.**
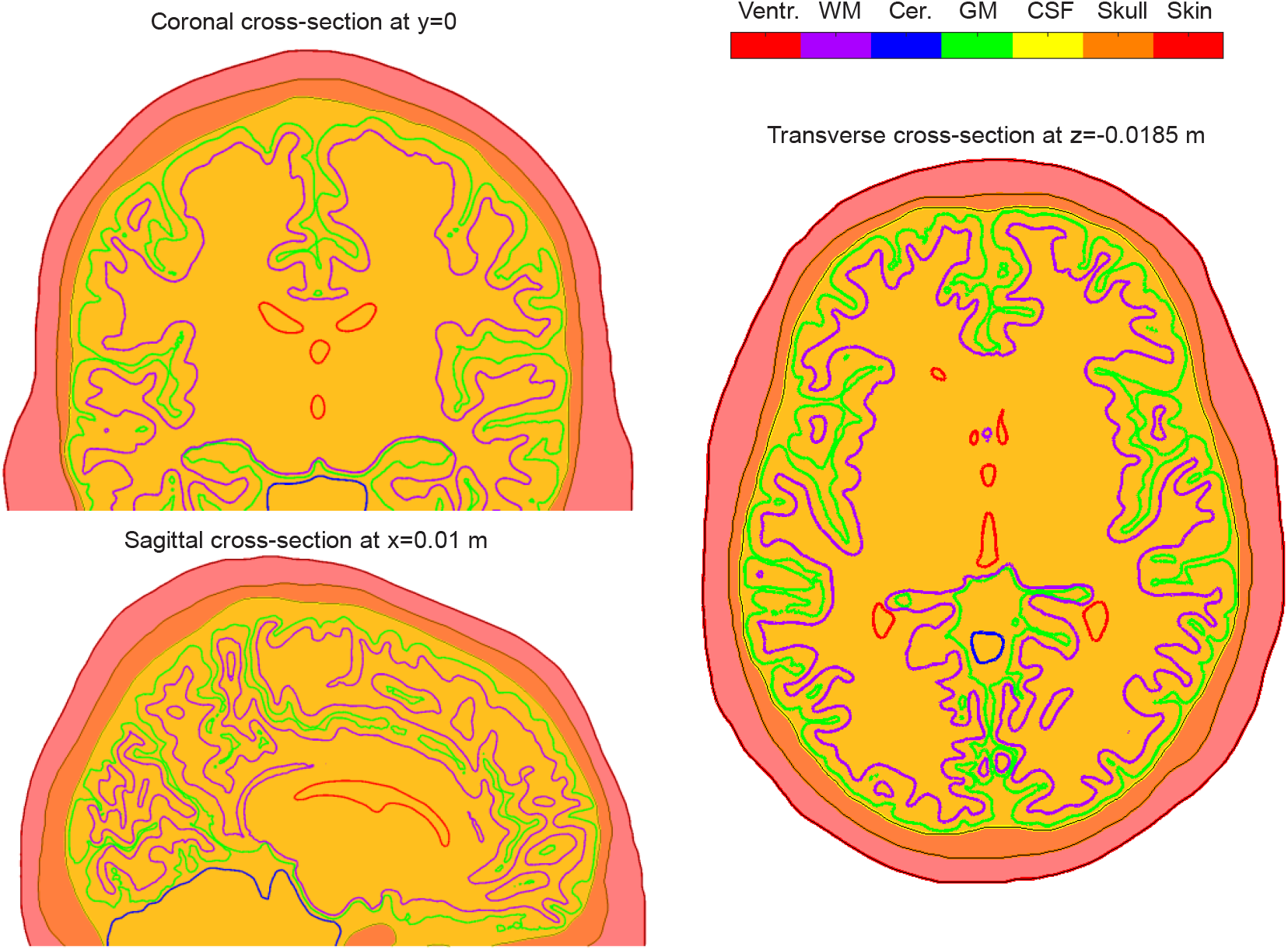
Three cross-section cuts obtained for subject # 110411 at *z* = −0.0185 *m, y* = 0, *x* = 0.01 *m*. Intracranial volume is marked yellow; skull volume is marked orange; and scalp volume is marked red.

For every head model we define a cortical layer as a volume within the GM shell but excluding WM volume. A dense point cloud within the cortical layer is additionally generated and used to estimate normal and tangential fields specifically in the cortical layer.

#### Precomputed global BEM-FMM parameters for the head model

The BEM-FMM algorithm needs to accurately compute electrostatic double surface field integrals for neighbor triangular facets *t_i_, t_j_* in the form:

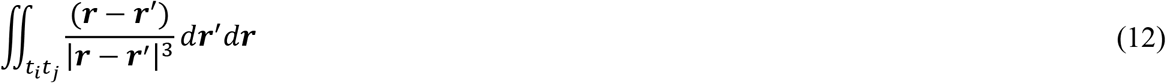

For each head model, these integrals are precomputed and saved. Their number could potentially be large. The safe default value is given by twelve neighbor integrals for every triangle *t_i_* excluding the self-integral, which is exactly zero. This value could be lowered to four if the corresponding file size appears too large. The typical file sizes to save the integrals are approximately 250 Mbytes.

### 2.3. Coil models

#### Coil CAD model

The base of a coil model is the conductor (or wire) centerline. The centerline of a conductor (or a multiple thereof) is created in MATLAB as set of 3D nodes distributed along a curve (circle, ellipse, spiral, etc.). After the centerline has been created, the CAD model for the coil conductor(s) is constructed by

i. Automatic creation of an unstructured triangular mesh for conductor’s cross-section similar to that shown in Fig. 2a,b. This mesh could have an arbitrary density and is either rectangular or elliptical in shape;
ii. Sweeping this cross-section mesh along curve’s centerline and creating a structured triangular surface mesh for the conductor’s side surface. The cross-section remains perpendicular to conductor’s centerline;
iii. Creating start/end caps for non-closed conductors and;
iv. Joining the meshes if necessary and eliminating duplicated nodes.

**Fig. 2.**
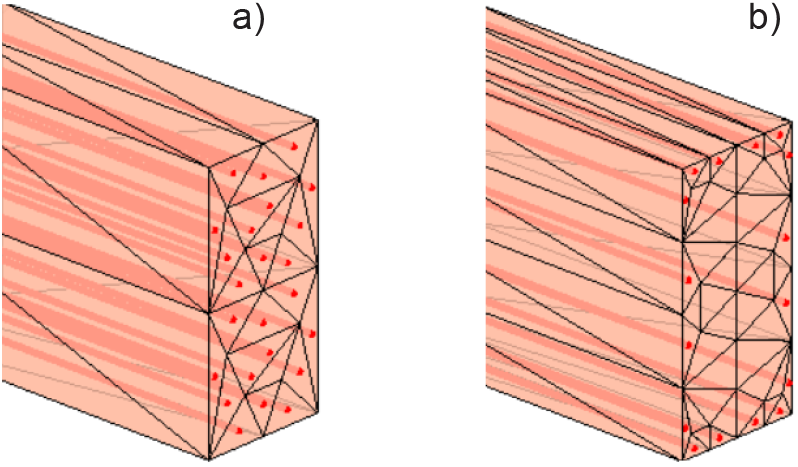
Filaments of current (red) inside conductor’s volume and conductor’s surface CAD model following cross-section triangulation. a) Modeling uniform current distribution through a conductor’s cross-section (Litz wire); b) Modeling the skin effect (a solid copper conductor at a higher frequency).

Fig. 3 shows simplified coil models created in this way. Fig. 3a is a simplified MRi-B91 TMS-MRI coil model (MagVenture, Denmark) with elliptical conductors of a rectangular cross-section; Fig. 3b is a simplified MagPro C-B60 coil model (MagVenture, Denmark); Fig. 3c is a double shape-eight spiral coil with an elliptical cross-section and two bootstrapped interconnections; Fig. 3d is a simplified Cool-40 Rat small animal coil model (MagVenture, Denmark); Fig. 3e is a three-axis multichannel TMS coil array radiator (Navarro de Lara et al, 2018).

**Fig. 3.**
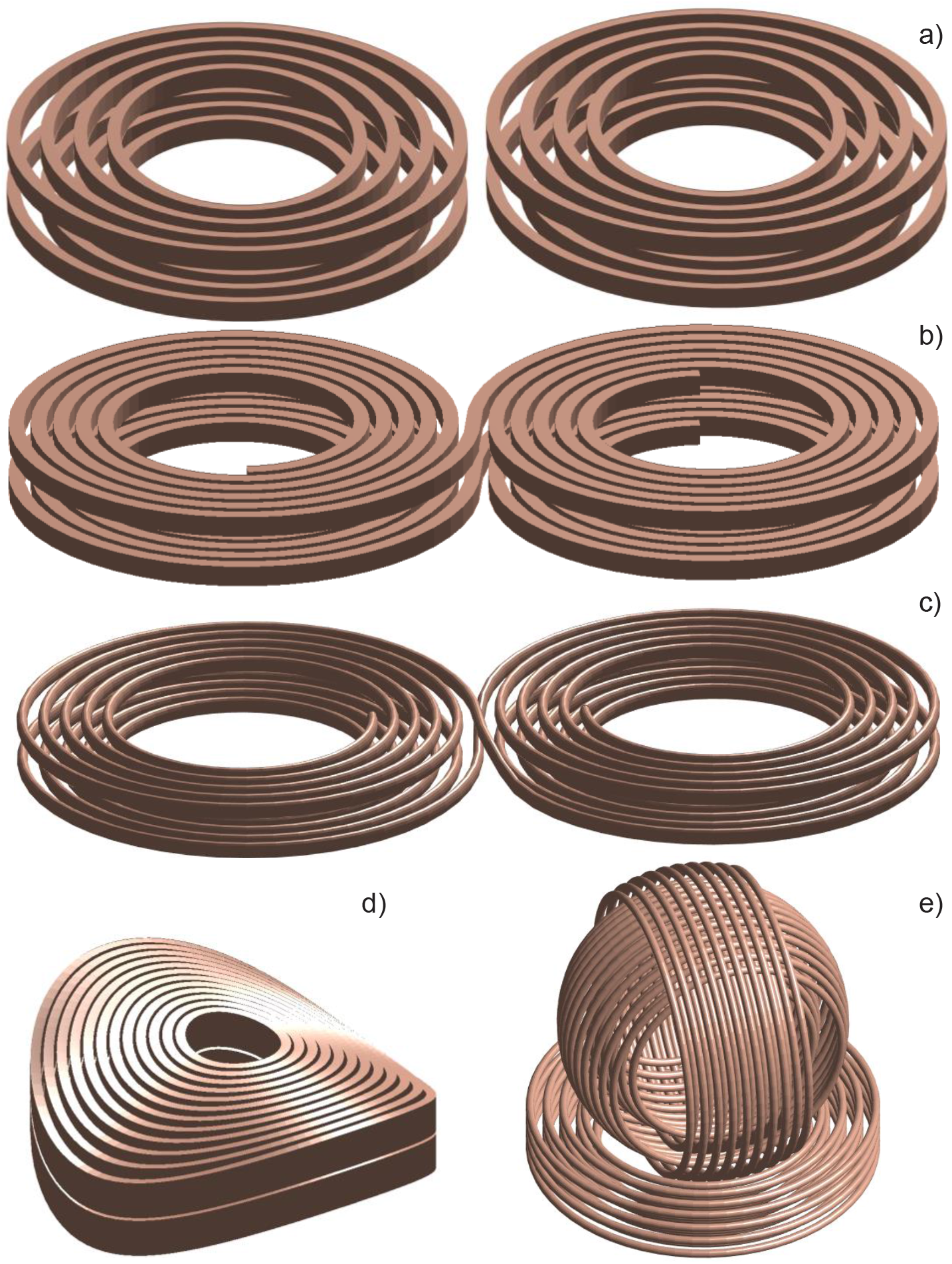
Some solid CAD models created using the MATLAB-based coil geometry generator. Fig. 3a is a simplified MRi-B91 TMS-MRI coil model (MagVenture, Denmark) with elliptical conductors of a rectangular cross-section; Fig. 3b is a simplified MagPro C-B60 coil model (MagVenture, Denmark); Fig. 3c is a generic double shape-eight spiral coil model with an elliptical cross-section and two bootstrapped interconnections; Fig. 3d is a simplified Cool-40 Rat small animal coil model (MagVenture, Denmark); Fig. 3e is a three-axis multichannel TMS coil array radiator (Navarro de Lara et al, 2018).

Once created in MATLAB, the CAD coil model may be exported to any other computational package in STL format. Note that coil models created in this way do not yet define the normal vectors of surface triangles. Therefore, after converting to STL format, a mesh healing operation is required; ANSYS Electronics Desktop and other packages implement this step.

#### Computational conductor model

The volumetric current flow is modeled as a set of infinity-thin current filaments or segments. The current filaments (shown red in Fig. 2a,b) are defined as short straight lines joining centroids of triangles of the cross-sectional mesh, which is replicated along the conductor’s centerline as many times as required. The cross-section is always perpendicular to the conductor’s centerline; the filaments are therefore parallel to the centerline of the conductor. The underlying computational wire grid is located strictly inside the solid conductor as illustrated in Fig. 2a,b.

The number of filaments passing through conductor’s cross-section can be made arbitrarily large. When a time-varying electric current flows in a volumetric conductor, two cases are possible: a) Litz-wire conductor with uniform current flow through its cross-section or b) current flow in a thin skin layer close to the surface. In the first case, the current filaments are uniformly distributed over the conductor’s cross-section as shown in Fig. 2a. In the second case, the current filaments are distributed close to its surface as shown in Fig 2b. Each filament carries current proportional to the area of the base triangle; the total conductor current is 1 A. Once the current filaments are internally defined, the primary field computations follow Eqs. (5), (8), and (9).

### 2.4 Software organization

Fig. 4 shows the software organization chart of the toolkit along with the major computation blocks (1 through 6). The blocks using the FMM computational algorithm, in some way or another, are marked grey. These program blocks do not have to be executed sequentially. They are described in more detail in Appendix A along with the program flow.

**Fig. 4.**
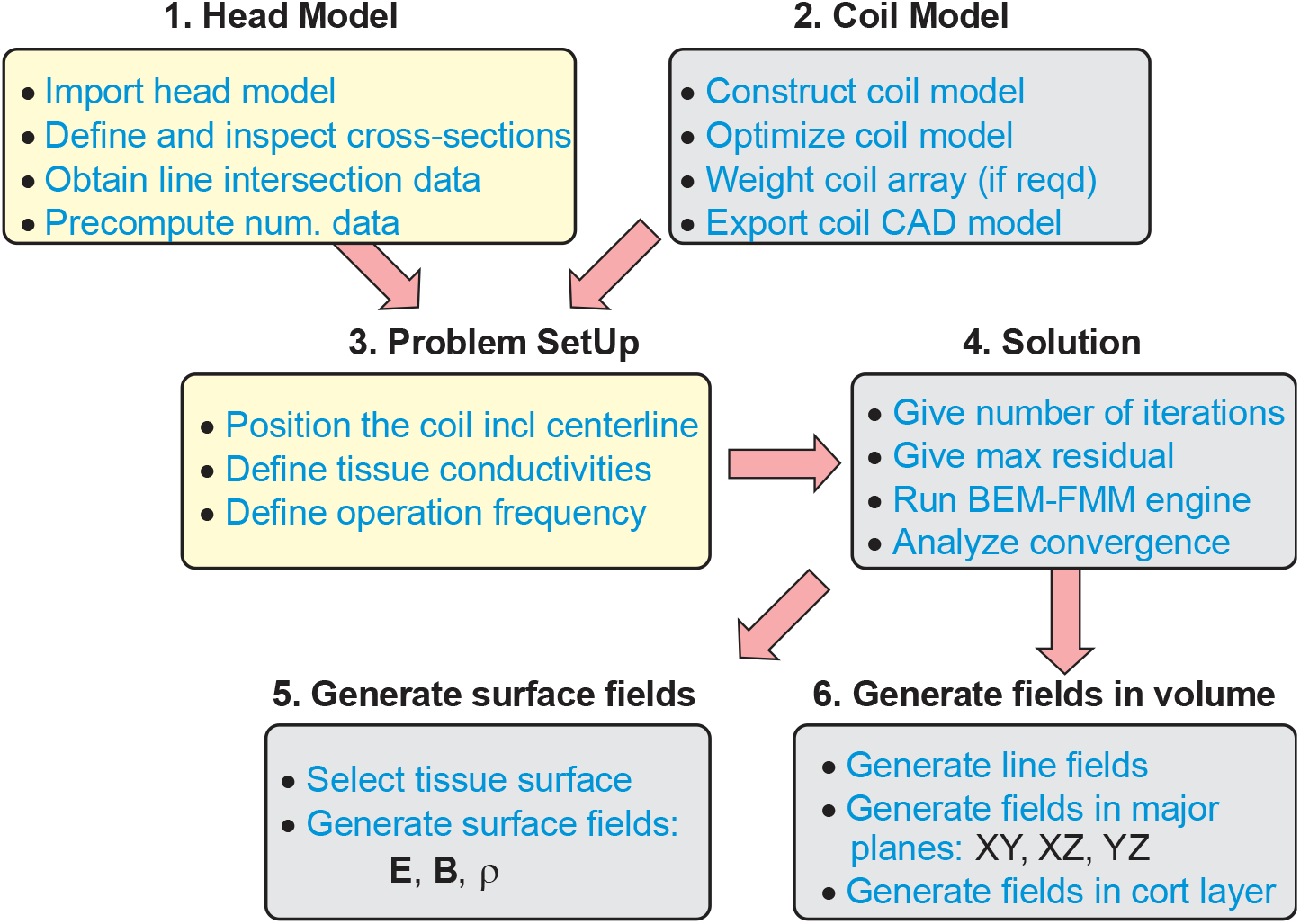
Software organization chart. Major computational blocks are labeled 1 through 6.

## 3. Results

### 3.1 FMM-based coil modeling in free space

The toolkit includes editable scripts for ten pre-constructed coil models, some of which are shown in Fig. 3. Other coil models or coil arrays can be constructed in a similar way, by defining the conductor centerline(s) and then automated cross-section sweeping. Of special note is the three-axis coil in Fig. 3e, which is in fact an array of three independent radiators with three independent excitations.

Coil modeling is done by computing and plotting respective electric and magnetic fields along a line or in a plane. Operating frequency (CW excitation is assumed) and conductor current must be specified. Fig. 5 shows typical output field results. Table 1 (Makarov et al., 2019) reports timing results for ***E***^*p*^ and ***B***^*p*^ computations. Here and below in the text, computations are performed using an Intel Xeon E5-2698 v4 CPU 2.2 GHz server, Windows Server 2016 operating system, and MATLAB 2018a.

**Fig. 5.**
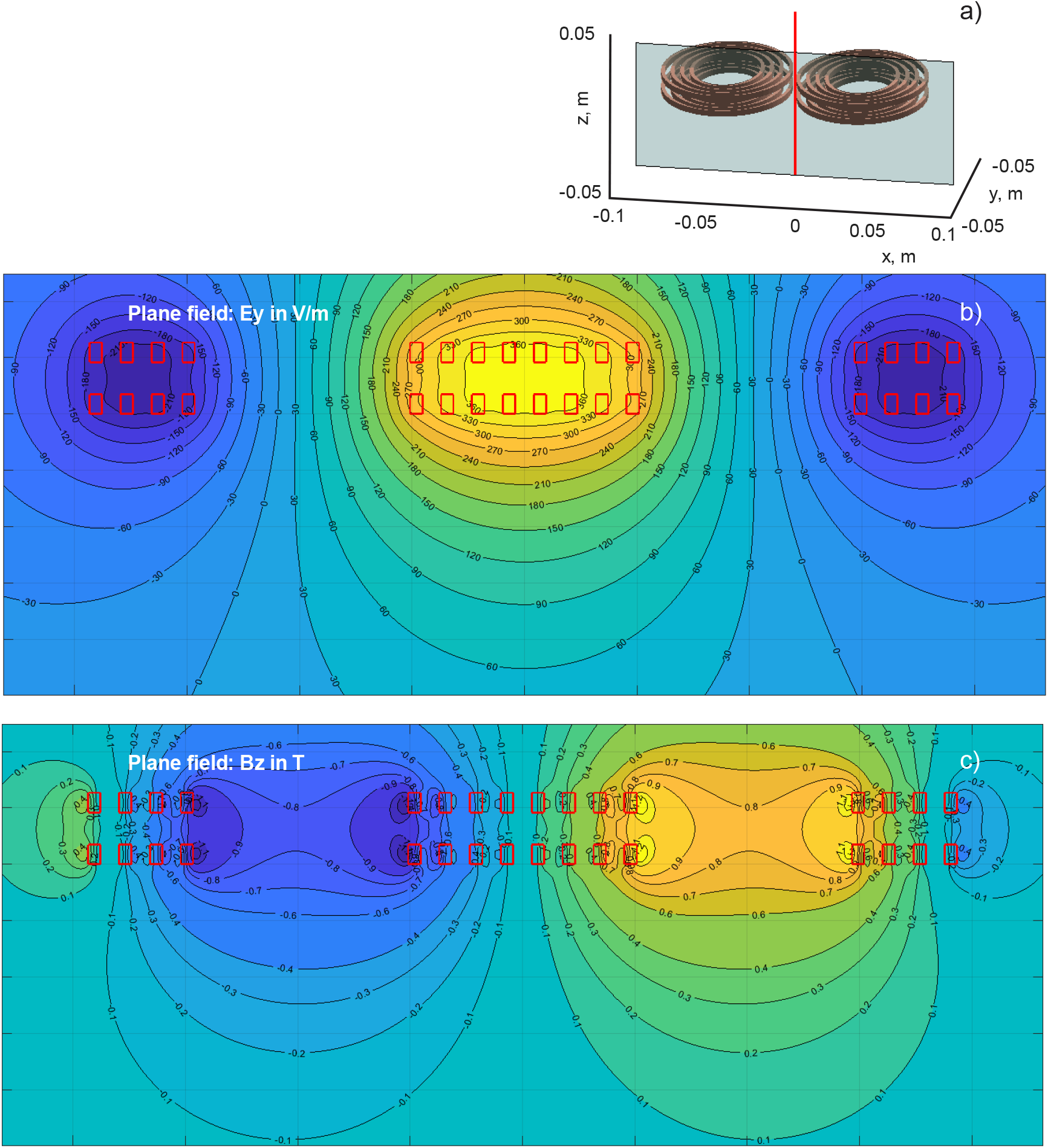
a) Observation plane geometry for MRi-B91 TMS-MRI coil model. b, c) Electric and magnetic fields in the plane with 0.25 M observation points. Conductor cross-section is marked in red. Conductor current is 5 kA and CW frequency is 3 kHz.

**Table 1.** Timing results for ***E***^*p*^ computations. An accurate coil model with 150,000 elementary current dipoles is considered.

| Observation points | Wire grid generation time, sec | FMM execution time, sec | Vectorized MATLAB loop, sec |
| --- | --- | --- | --- |
| 100×100 | 0.32 | 0.44 | 92 |
| 200×200 | 0.32 | 0.47 | 394 |
| 300×300 | 0.32 | 0.57 | 892 |
| 500×500 | 0.32 | 0.93 | 2503 |

As an example, a conical-shape coil with 50 single coaxial loops of a circular cross-section has been considered. Each of the loops is divided into 100 straight filaments. We assume a Litz-wire conductor and introduce 30 interpolation points nearly uniformly distributed over conductor’s cross-section. This results in a coil model that contains 150,000 elementary current dipoles in total.

We now keep this coil geometry fixed but vary the number of observation points in a coronal plane whose size is twice the coil size as per Tables 1 and 2, respectively. In these tables, we present mesh generation time for the wire grid, FMM execution times, and execution times for a simple yet vectorized MATLAB code that directly computes Eq. (5) and Eq. (7) for the required set of observation points. All results have been averaged over several runs.

**Table 2.** Timing results for ***B***^*p*^ computations. An accurate coil model with 150,000 elementary current dipoles is considered.

| Observation points | Wire grid generation time, sec | FMM execution time, sec | Vectorized MATLAB loop, sec |
| --- | --- | --- | --- |
| 100×100 | 0.31 | 0.49 | 242 |
| 200×200 | 0.31 | 0.52 | 968 |
| 300×300 | 0.31 | 0.62 | 2250 |
| 500×500 | 0.31 | 0.99 | 6320 |

By analyzing the representative FMM computations and the underlying wire model generation times from Tables 1 and 2, we observe that coil evaluation can generally be performed in less than 1 sec on an ordinary server. On the other hand, the direct vectorized MATLAB loop would run 200-6,000 times slower.

We also observe that the FMM code indicates only a very modest increase in time for the reported size of the coil model and the sizes of the observation domain. No specific effort to parallelize the algorithm has yet been made. However, MATLAB automatically performs multithreading pertinent to linear algebra operations available in LAPACK (Linear Algebra PACKage) and some level-3 BLAS (Basic Linear Algebra Subprograms) matrix operations, allowing them to execute faster on multicore-enabled machines.

With regard to coil optimization, parametric loops versus coil geometry parameters could straightforwardly be organized and executed relatively fast. Reference Makarov et al., 2019 discusses corresponding examples. Additionally, an example pertinent to steering in a multi-channel coil array is also discussed.

### 3.2. Electric field computations within the head

#### Computational times

For head models with ~0.9 M facets described above, the main iterative algorithm runs in approximately 90 sec (4.5 sec per iteration for 20 iterations). Restoration of surface electric fields (just inside or outside a conducting boundary) requires one step of the iterative solution, i.e. 4-5 sec. Restoration of line fields is also fast. However, detailed field computations with a fine resolution (0.3-0.4 mm) in a plane crossing many conductivity boundaries as well as field computations in a cortical volume require 30-40 sec. This is because the contributions of surface charges on nearby triangles need to be accurately averaged for every observation point in question, similar to Eq. (11). The run time data are for an Intel Xeon E5-2698 v4 CPU (2.20 GHz) server, 256 Gb RAM, MATLAB 2018a.

#### Computational example

In the following computation example, we will consider Connectome head model # 110411 and MRi-B91 TMS-MRI coil model targeting the area of the primary motor cortex and located above the precentral gyrus of the right hemisphere. The tissue conductivity values are those of Engwer et al., 2017 and Piastra et al., 2018. Conductor current is 5 kA and CW frequency is 3 kHz. Fig. 6a shows the model, the coil position above the skin surface, the coil centerline (red) which starts at the coil bottom, as well as positions and sizes of three principal observation planes. All three planes pass through the point which is an intersection of the gray matter shell with the coil centerline.

**Fig.6.**
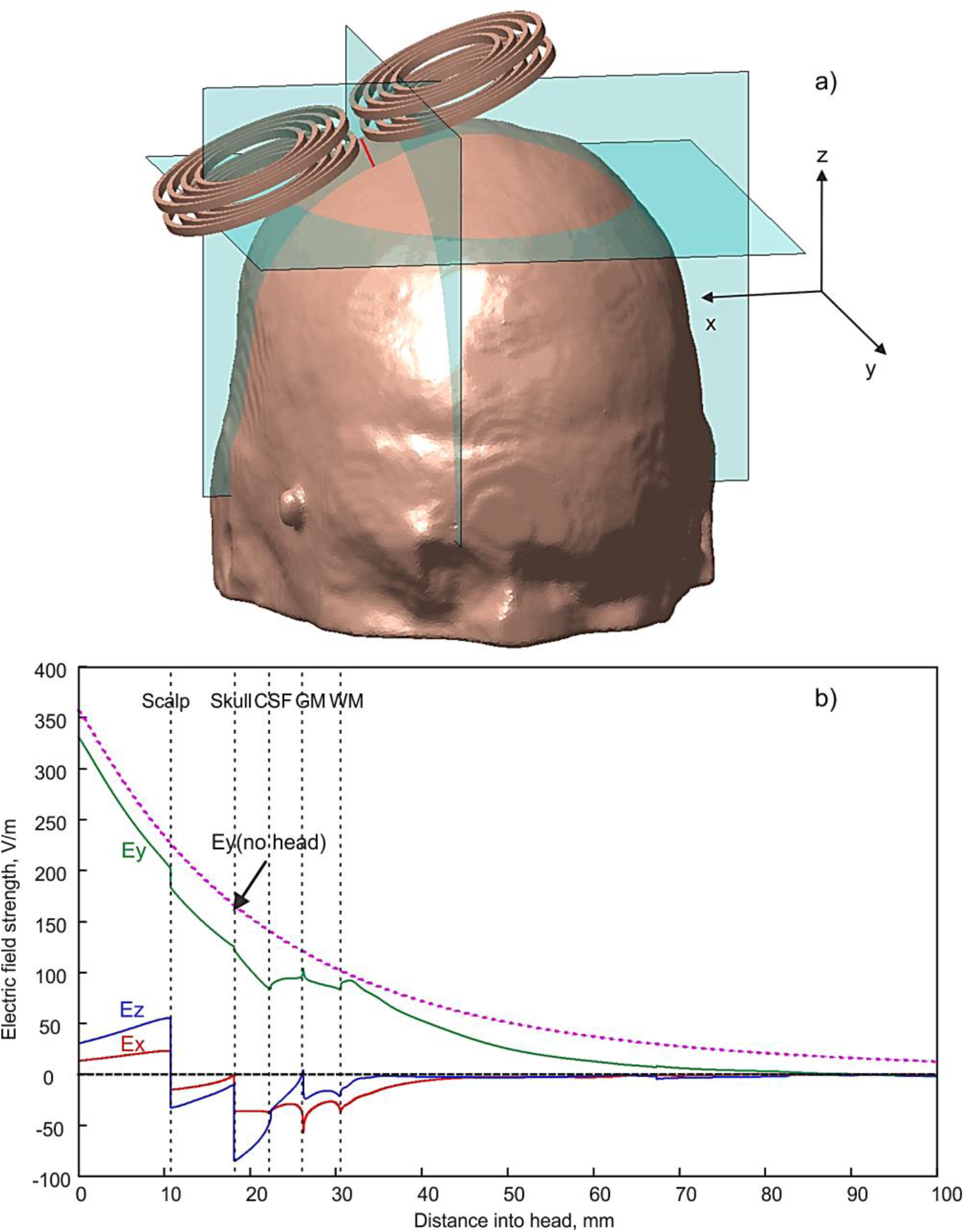
a) – Computation geometry, position of the observation line (coil centerline), and positions of three observation planes. b) Electric field distribution along the coil centerline.

#### Fields along a line

Fig. 6b shows three Cartesian components of the electric field computed along the coil centerline. Any other line may be considered as well. The prominent field discontinuity at tissue interfaces is due to surface charges residing at the conductor boundaries. This discontinuity is precisely tracked with our computational method. Comparison of line fields with the commercial FEM software ANSYS Maxwell, which uses adaptive mesh refinement, has been performed in (Htet et al., 2019). There, ten head models from the Population Head Model Repository (Lee et al., 2016; Lee et al., 2018; The Population Head Model Repository, 2017) have been tested and a very good agreement between the two approaches was established.

#### Surface fields just inside/outside brain compartments (conductivity boundaries)

For any model shell (skin, skull, CSF, GM, WM, etc.) the fields and surface charge distribution can be extracted. The magnetic field is simply the continuous coil’s field in free space, ***B***^*t*^(***r***) = ***B***^*p*^(*r*), since the secondary field of eddy currents is ignored in TMS studies. The total electric field, however, is discontinuous at interfaces. When approaching an interface from outside/inside it is given by

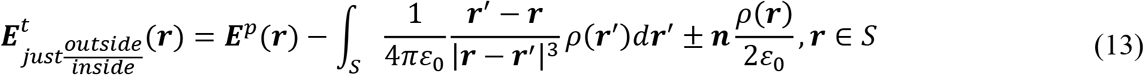

where the notations have been explained in Section 2.1. Since the charge distribution is already known from the BEM-FMM solution, the integral on the right-hand side of Eq. (13) is computed using the FMM (with accurate neighbor integral contributions). Then, the electric field just inside or outside the interface is obtained by simply choosing the proper sign in front of the last term on the right-hand side of Eq. (13).

Figs. 7a-c show surface electric fields just inside the gray matter shell, i.e. at the top of the cortical layer. In Fig. 7a, the magnitude of total electric field is shown. Fig. 7b presents the magnitude of the continuous field component tangential to the surface and Fig. 7c is the most important discontinuous signed normal electric field. The color scale expands from 0 to 160 V/m, 0 to 120 V/m, and –150 to 150 V/m, respectively.

**Fig. 7.**
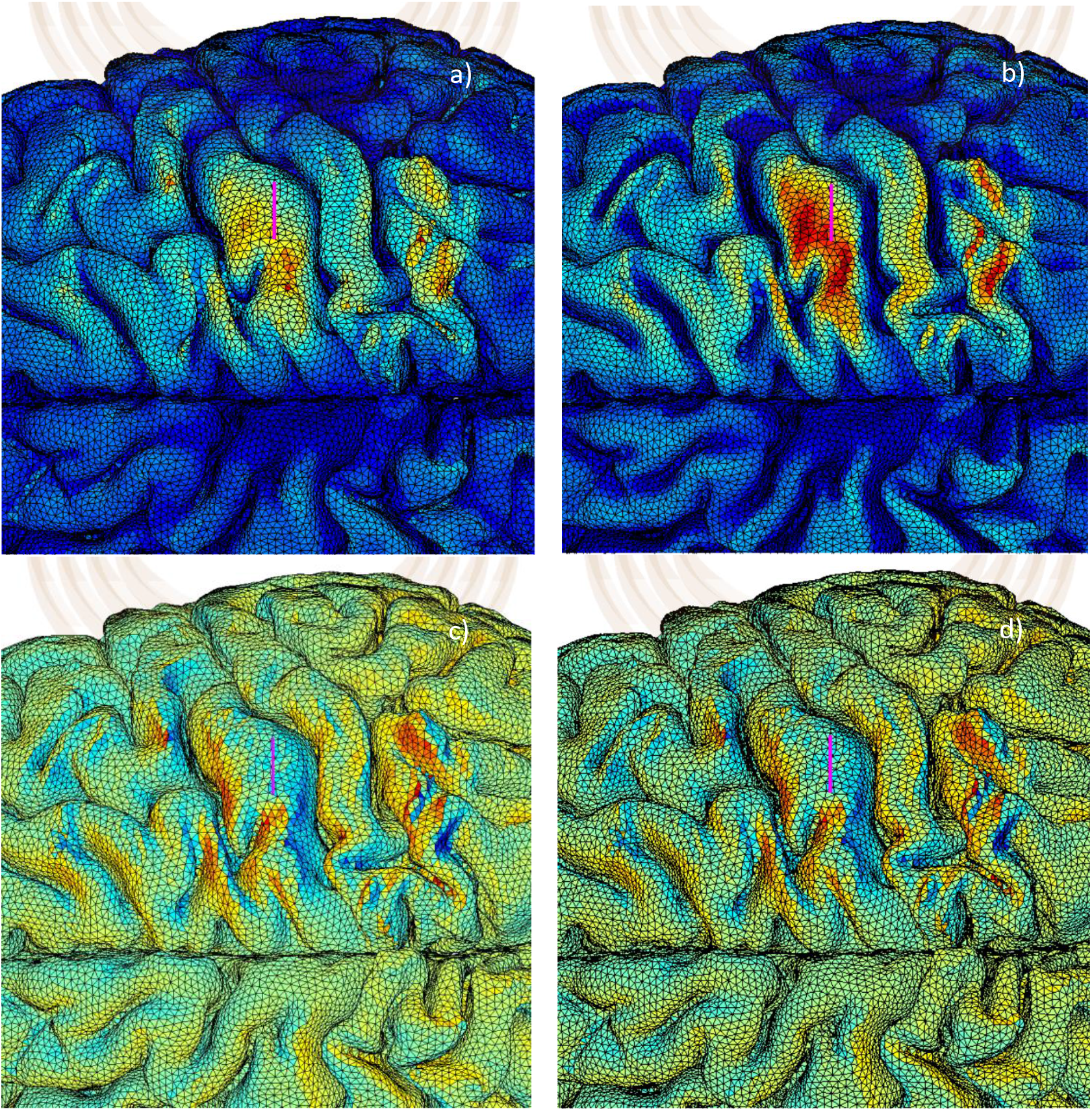
Surface electric fields just inside the gray matter shell: the top level of the cortical layer. a) Magnitude of total electric field (0-160 V/m). b) Magnitude of tangential electric field (0-120 V/m). c) Signed normal electric field component (−150 to 150 V/m) in the direction of the inner normal vector, i.e. into the gray matter volume. d) Induced surface electric charge density on the gray matter surface; the color scale expands from −1 to +1×10^-9^ C/m^2^.

Total normal and tangential fields given in Fig. 7b,ca are defined in the form:

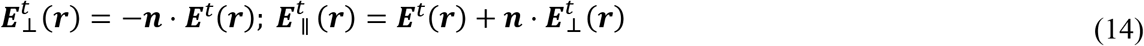

i.e. the signed normal field 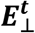 is considered positive if it is directed into the cortical layer. The total and tangential fields in Fig. 7a,b indicate rather familiar behavior referenced in many literature sources. The normal field in Fig. 7c is more irregular. It is mostly concentrated inside certain sulci, where the dominant coil field component becomes perpendicular to the cortical surface. A sulcus that is perpendicular to the dominant field component thus becomes “polarized”.

Quite remarkably, the inner normal field component in Fig. 7c almost perfectly correlates with the underlying induced surface charge density distribution on the gray matter surface shown in Fig 7d. There, the color scale expands from −1 to +1×10^−9^ C/m^2^. The correlation measure is the relative error, Δ***E***, given by

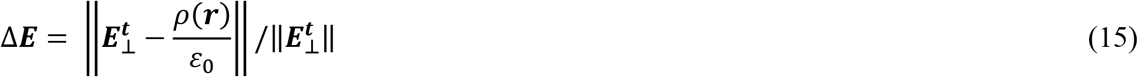

The error value (15) for the present example for the entire gray matter shell is 5.3%. This error value decreases further when only the area under the coil is considered. Such an observation might be significant for real-time TMS field computations since computing and storing the charge distributions is far more efficient than computing and storing the field distributions. This question, however, is beyond the scope of the present study.

#### Fields in principal planes

Those computations require more time since the potential integrals are no longer precomputed and must be calculated in real time. Fig. 8a illustrates the electric field distribution for the y-component of the total electric field (the component that is directed out of figure plane) in the coronal plane from Fig. 6 with a grid resolution of 0.4×0.3 mm.

**Fig. 8.**
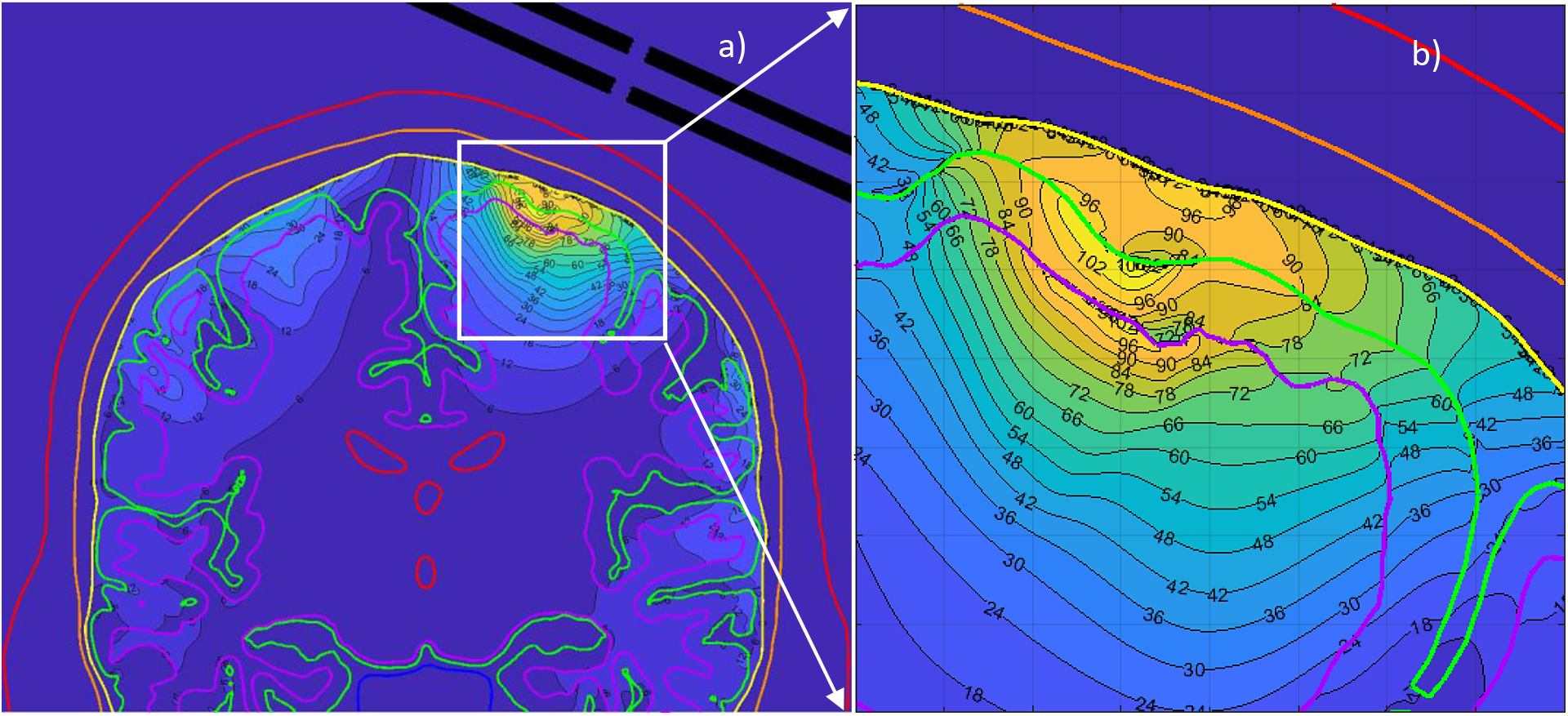
Y-component of the total electric field (directed out of figure plane) in the coronal plane from Fig. 6 with a grid resolution of 0.4×0.3 mm (0.25 M observation points). The color scale expands from 0 to 105 V/m. a) Field in the entire plane. b) Field in a zoomed in cortical area with the size of 40×40 mm.

The y-component is simultaneously the dominant primary field component of the coil. Only the field within brain compartments excluding scalp and skull is displayed in Fig. 8. The color scale expands for 0 to 105 V/m.

By observing Fig. 8, it might appear at first sight that the field component of interest is well localized just underneath the coil. However, this observation is only valid for this particular plane and for this particular (primarily tangential to the cortex) field component.

#### Volumetric fields within the cortical volume

We define the cortical layer as a volume within the GM shell but excluding the WM volume. A point cloud within the cortical layer is automatically generated and used to estimate normal and tangential fields specifically in the cortical layer. This point cloud contains at least five inner observation points across the layer, which are uniformly spaced along the normal direction of every triangle of the GM shell. Both normal and tangential field components are evaluated at every such observation point and respective “hot spots” within the cortical layer are identified and labeled with a small prominent blue ball.

Fig. 9a shows examples of hot spots where the magnitude of the total normal field component is greater than or equal to 90% of the maximum total normal field in the cortex; Fig. 9b shows a similar result but for the 80% threshold. The underlying white matter shell is also shown. One observes that the hot spots for the peak fields may be very well localized in space as long as the required field is 80% or higher of the maximum normal field (for this case the maximum normal field is approximately 140 V/m). In this sense, the focality of the coil could be well established for a specific position/subject. However, these hot spots may occur not directly beneath the coil as expected, but at a rather significant distance from the coil centerline as seen in Fig. 9a,b. This observation underscores the value of numerical modeling.

**Fig. 9.**
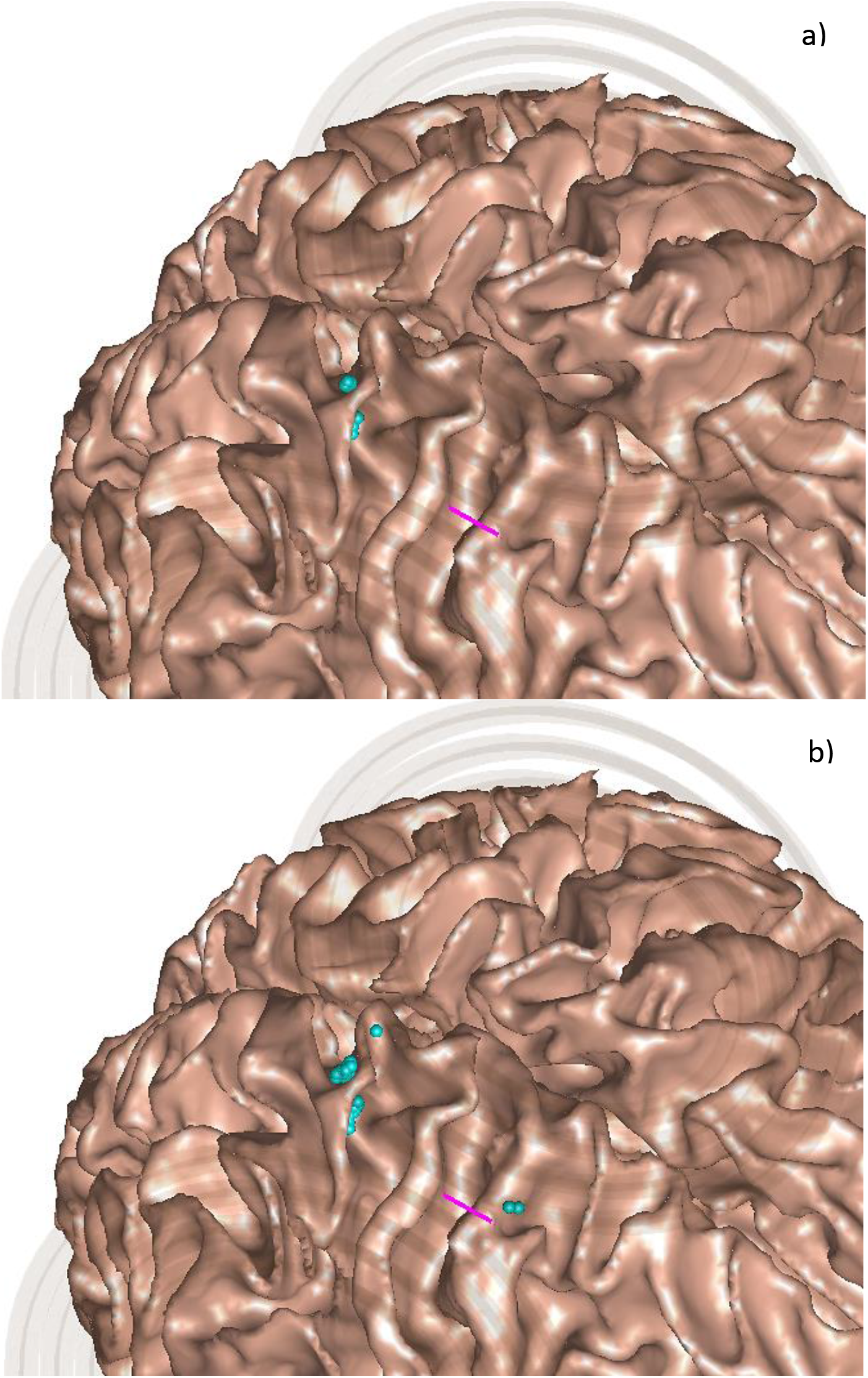
Hot spots – observation points within the cortical volume where the magnitude of the total normal field component is greater than or equal to 90% (a) or 80% (b) of the maximum total normal field in the cortex. The underlying white matter shell is shown.

## 4. Discussion and Conclusions

In the present study, we have described the performance and capabilities of a software toolkit for high-resolution TMS modeling. The toolkit employs the boundary element fast multipole method with accurate field evaluation close to conducting interfaces. It operates entirely within the MATLAB platform and is designed for a broad, medically-oriented audience without specialized programming skills. To enable fast, subject-specific modeling, we have constructed a simulation capability compatible with automated human head CAD model generation tools such as SimNIBS or FreeSurfer. The toolkit is augmented with the collection of 16 head models and with a coil geometry generator script.

Both coil design in free space as well as efficacy and focality of TMS for a specific subject could be modeled by the package, including optional user-defined MATLAB parametric loops. Detailed coil models, presented in both CAD and wire formats, may include several hundred thousand elementary current elements and observation spaces with approximately 1 M field points; the corresponding computational times are on the order of 1 sec. Detailed head models with approximately 1 M triangular facets and a mesh resolution of 0.6 points per mm^2^ are simulated in approximately 1.5 min. Further reduction of the computational time is foreseen by using real-valued FMM arithmetic.

The fields within the head are examined along a line, just inside or outside a brain compartment surface, in three principal planes, or in the entire cortical volume. Volumetric field computations with a large number of observation points close to charged boundaries may require up to 1 min.

Although the accuracy of the present numerical approach has already been justified, an additional error test may be needed in a specific case. To do so, a refined set of head models has been included into the toolkit with 3.5 M triangular facets on average and with a mesh resolution of 1.2 points per mm^2^. These were obtained after barycentric oversampling (4×1) and subsequent surface-preserving Laplacian smoothing (Vollmer et al., 1999) of the base models. Those models may be characterized by a better solution convergence; they run in approximately 200-250 sec.

In conclusion, we anticipate that the described TMS modeling toolkit will be a useful academic research tool. The authors would welcome user feedback and look forward to further improving software performance and capabilities.

## Acknowledgements

The authors wish to thank Dr. Leslie Greengard of the Courant Institute of Mathematical Sciences, New York, NY for multiple insightful discussions and support in our utilization of the FMM library. This work has been partially supported by the National Institutes of Health under award numbers R43AR071220, R00EB015445, NINDS R44NS090894, and R01MH111829.

## Code availability

The code will be made available through

i. MATLAB Central. 2019 Onl.: https://www.mathworks.com/matlabcentral/and;
ii. Athinoula A. Martinos Center for Biomedical Imaging, Massachusetts General Hospital. 2019 Onl.: https://www.nmr.mgh.harvard.edu/research/software

These links are not yet active. To obtain the toolkit code, please send a short note to the first author.

## Appendix A. Concise User’s Manual

### A1. Low-level organization chart

The toolkit contains a number of MATLAB scripts organized within three subfolders – `Model, Coil`, and `Engine` – and a number of scripts located in the main folder, as shown in Fig. A1. The subfolder Model deals with a particular head model definition and precomputations. The subfolder Coil handles coil definition, construction, and testing/optimization. The subfolder Engine contains computational scripts which should not be changed unless necessary. The scripts from the main folder perform major numerical operations and output the fields.

**Fig. A1.**
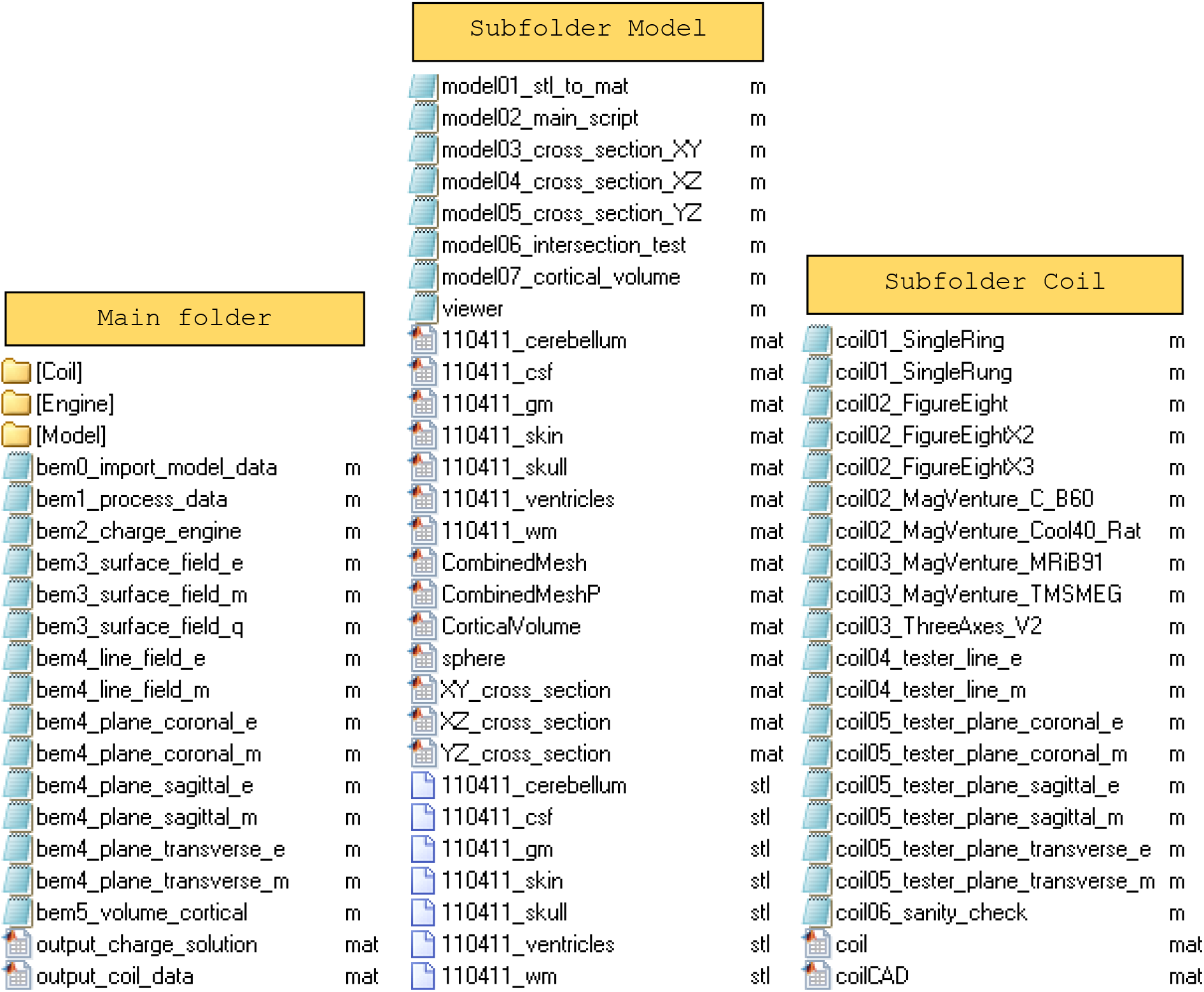
Low-level organization chart of the toolkit.

### A2. Computational sequence of the main folder

The scripts of the main folder could be executed at any time since all data have already been precomputed. The first two scripts must be executed sequentially, i.e. `bem0_import_model_data.m` first and then `bem1_process_data.m` next. The first script imports model data into the MATLAB workspace and sets the MATLAB path. The second script initializes (i) operation frequency and coil current; (ii) tissue conductivities; (iii) coil position above the head by proper rotation and translation; (iv) coil centerline or another observation line and; (v) displays combined head-coil geometry as shown in Fig. 6a of the main text. Coil positon adjustment may be performed by running the script `bem1_process_data` multiple times. When performing mathematical (e.g. coil rotation) and graphical operations, this script calls a number of functions from subfolder `Engine`.

The next script to be executed is `bem2_charge_engine.m`. This script (i) computes the primary field of the coil on every head interface using FMM; (ii) computes the iterative solution of Eq. (2) of the main text for the induced surface charge density using the FMM, precomputed near-field potential integrals, and MATLAB GMRES, and; (iii) displays iteration time for every iteration step in the MATLAB command window and then plots the entire convergence history when completed. Since no charge conservation law is explicitly enforced in the current version, the resulting charge error (typically below 0.5%) is also computed and displayed. The maximum permissible number of iterations and the targeted relative residual are defined at the beginning of the script. It is not recommended to change these default values. The execution time is approximately 1.5 min. When performing mathematical (FMM) operations, this script calls original and derived FMM functions from subfolder `Engine`.

Upon completion of this step, any of the field computational scripts may be executed, in any sequence. The script `bem3_surface_field_e` computes (via the FMM) and displays generally discontinuous electric field just inside or outside any head compartment, as shown in Figs. 7a,b,c of the main text. Normal (signed), tangential (magnitude), and total (magnitude) field components may be plotted. The script `bem3_surface_field_m` performs a similar operation for the magnetic field which remains continuous across the boundaries. The script `bem3_surface_field_q` displays induced surface charge density as shown in Fig. 7d of the main text.

Scripts `bem4_line_field_e/m.m` compute the three components of the electric/magnetic field along the coil centerline or, alternatively along any arbitrary line using the FMM with accurately averaged contributions of the surface charge density on neighbor triangles. An example is given in Fig. 8 of the main text. There and in the following scripts, parameter *R* characterizes dimensionless (vs. average edge length) radius of the integration sphere with the center at the observation point within which precise integration over triangular facets is performed.

The following scripts starting with `bem4_plane_coronal_e.m, bem4_plane_coronal_m.m` etc. output electric or magnetic field (any of its Cartesian components or a magnitude) in three principal planes. Accuracy of the electric field computation in the vicinity of the compartment boundaries is again controlled by parameter *R* – dimensionless radius of the precise-integration sphere. Finally, the script `bem5_volume_cortical.m` displays domains of the maximum normal or tangential electric field within the cortical layer between gray matter and white matter shells. A representative example is given in Fig. 9 of the main text. When performing mathematical (FMM) and graphical operations, all of these scripts call a number of functions from subfolder `Engine`.

### A3. Computational sequence of subfolder Model

The head model files are located in the dedicated folder `Model` with contents shown in Fig. A1. The primary set are *.stl (stereolithography) files for every individual brain compartment in the form of a faceted shell with triangular facets and the respective outer normal vectors. The number of shells may be arbitrary. Every stl file, either binary or ASCII, is then converted to an equivalent MATLAB data file seen in Fig. A1 with arrays of vertices, *P*, facets, *t*, and normal vectors, ***n***, using the script `model01_stl_to_mat`. Every MATLAB data file can inspected and visualized using the `viewer` function from the same subfolder as shown in Fig. 1a of the main text. Next, the combined mesh for the entire head is created by appending individual meshes. The combined mesh is stored in the MATLAB data file `CombinedMesh.mat`. This is done by running the script `model02_main_script.m`. There is also an additional and bigger data file `CombinedHeadP.mat` generated in the same folder, which contains precomputed double surface electrostatic integrals over triangles from Eq. (11) necessary for accurate BEM-FMM simulations. Computations of integrals are done in parallel and using the default number of 24 cores. The `parfor` loop of MATLAB may be adjusted or removed depending on the computer configuration.

The present package is augmented with 16 head models for 16 Connectome subjects (101309, 110411, 117122, 120111, 122317, 122620, 124422, 128632, 130013, 131722, 138534, 149337, 149539, 151627, 160123, 198451). These models (and their refined versions) described in (Htet et al., 2018) could be downloaded independently from the MATLAB Central link (Collection of Sixteen High-Quality Human Head CAD Models. 2019).

The following three scripts are `model03_cross_section_XY.m, model03_cross_section_XZ.m`, and `model03_cross_section_YZ.m`. They compute tissue mesh intersections with three principal planes specified by the user. The intersection contour for every tissue is found and saved in the respective MATLAB data file. For skin, skull, and CSF shells, such a contour is given by a 2 manifold polygon, which could be filled. An example is given in Fig. 1b.

The script `model06_intersection_test.m` finds tissue mesh intersection points with an arbitrary line. These data may be useful for coil positioning. Finally, the script `model07_cortical_volume.m` creates and saves a point cloud between gray matter and white matter shells: the cortical layer. This point cloud contains five or more inner observation points across the layer, which are uniformly spaced along the normal direction of every triangle of the GM shell. It is used to compute hot spots of the electric field within the cortex as shown in Fig. 9 of the main text. When performing mathematical (potential integrals, mesh intersections) and graphical operations, these scripts call a number of functions from subfolder `Engine`.

### A4. Computational sequence of subfolder Coil

First, a coil model (both wire and CAD) is generated using a dedicated MATLAB script, with *one* script per coil. The bulk of the manual work here is the definition of the conductor’s centerline using either an analytical formula or a set of points in three dimensions. After that, the script automatically generates the volumetric computational wire grid coil model and the illustrative CAD coil model. The default coil axis is the *z*-axis.

The first task (wire model shown in Fig. 2 of the main text) is accomplished by the function `meshwire.m` of subfolder `Engine`. This function creates the wire mesh for an arbitrary *single* conductor. Either closed loops (`coil01_SingleRing.m`) or open conductors (`coil01_SingleRung.m`) may be considered. The computational wire grid coil model consists of straight, short, infinitely-thin current filaments or segments. The current filaments (shown in red in Fig. 2 of the main text) are defined as short straight lines joining centroids of triangles of the cross-sectional mesh, which is replicated along the conductor’s centerline as many times as required. The cross-section is always perpendicular to the conductor’s centerline; the filaments are therefore parallel to the centerline of the conductor. The result is structure `strcoil` with the following fields:

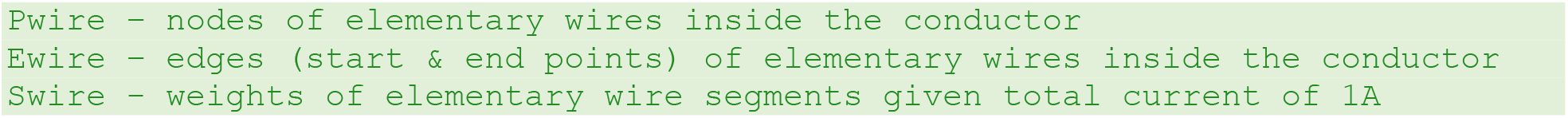

Weights are necessary to ensure that the total current through the conductor’s cross-section is 1 A.

Along with the function `meshwire.m`, the function `meshcoil.m` can be used. This function performs a similar task, but for a particular coil geometry consisting of a (large) number of concentric loops; it creates the wire coil model all at once. The input are intersection points of the loop centerlines with *xz*- and *yz* planes.

Either a Litz wire model (parameter `sk = 0`) or a skin-layer model (parameter `sk = 1`) can be considered in both functions `meshwire.m`, and `meshcoil.m`. In the former case, the current distribution across conductor’s gross-section is nearly uniform. In the latter case, the wire grid is situated close to the surface of the conductor. The density of the wire grid depends on cross-section triangulation; it is controlled by parameter `M` – the number of cross-section subdivisions. The grid resolution in the direction of the conductor centerline is controlled by the original centerline discretization.

A CAD model for the coil conductor is constructed using the function `meshsurface.m` of subfolder `Engine` and by creating a structured triangular surface mesh for the conductor’s side surface. This mesh is created based on the array of edges *e* for the conductor’s perimeter (which may either be elliptical or rectangular); the perimeter contour stays the same when moving along the centerline. At every discrete step along the conductor’s centerline (such steps are illustrated in Fig. A2), we add triangular facets in an ordered way. The corresponding MATLAB code accumulates side facets into array *t* as follows where *NE* is the number of edges (and nodes) for the conductor’s perimeter contour. At every step, we also add *NE* new nodes. Next, start/end caps for non-closed conductors are created and duplicated nodes are removed. As a result, the solid CAD model of a single coil conductor and the entire coil is created as shown in Fig. 3 of the main text.

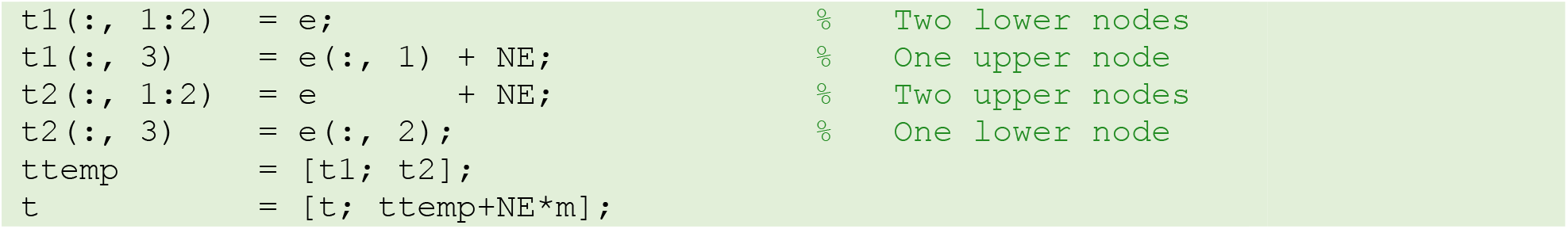

Note that the coil model does not yet define the normal vectors of triangular surface patches, nor does it properly orient the triangles. Therefore, after converting to STL format, a mesh healing operation is required, for example in ANSYS Electronics Desktop. Fig. A2 shows a more detailed concept of the combined wire/CAD model.

**Fig. A2.**
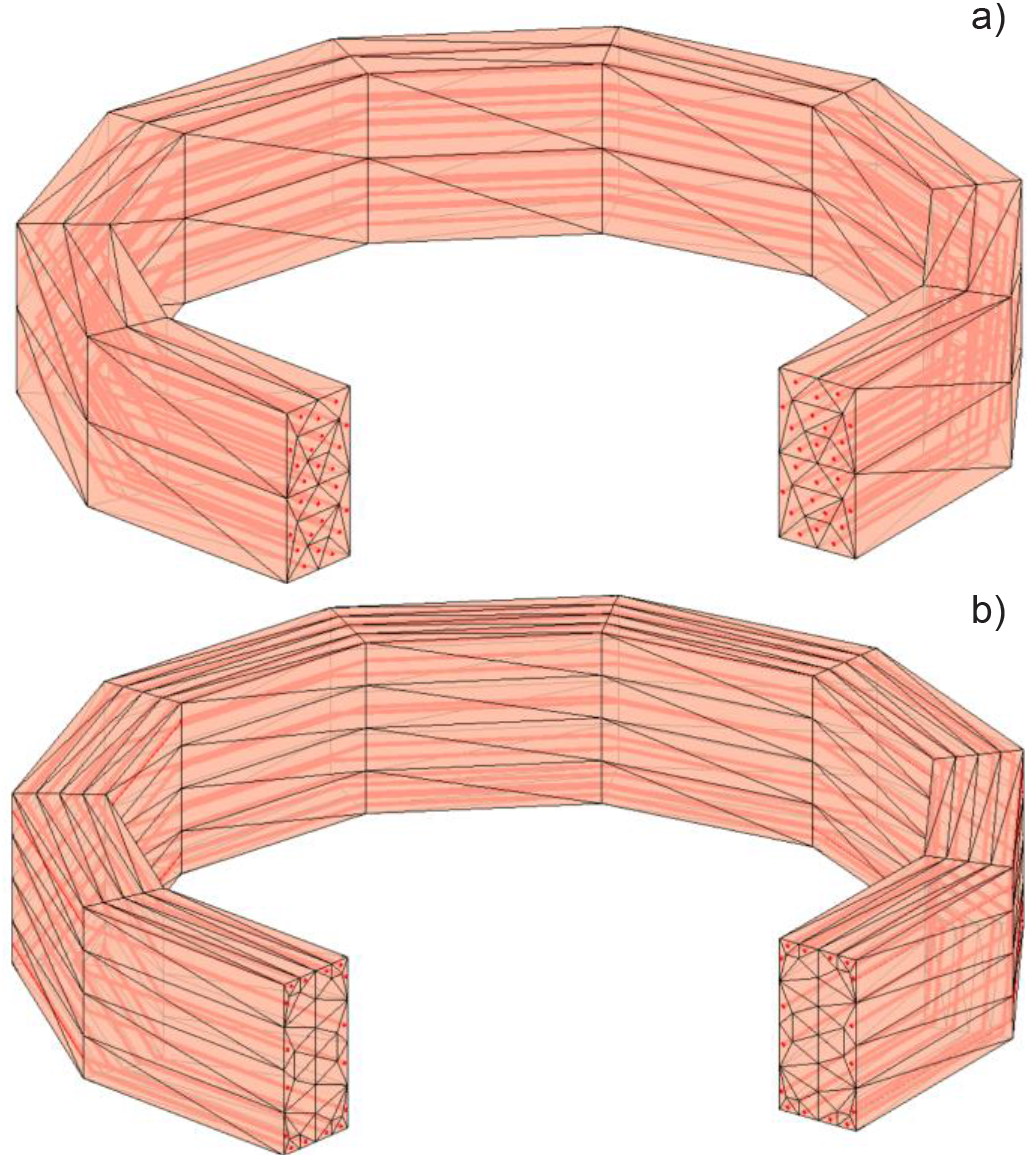
Filaments of current (red) within conductor’s surface CAD model. a) – Uniform current distribution (Litz wire); b) – modeling the skin effect (a solid conductor at a high frequency).

At present, the coil geometry modeler is restricted to predominantly flat or moderately bent conductor loops or nearly planar curves. H-coils (see, for example, Deng, etal., 2013) with sharp conductor bends in all three planes may be subject to further investigation, along with coil inductance calculations.

After the coil model has been created, line plots (`coil04_tester_line_e/m.m`) as well as high-resolution 2D contour plots (`coil05_tester_plane_coronal_e/m.m` etc.) for any component of the electric and/or magnetic field in the coronal, sagittal, and transverse planes are computed via the FMM, similar to the plots shown in Fig. 5 of the main text. When performing mathematical (mesh generation, FMM computations) and graphical operations, these scripts call a number of functions from subfolder `Engine`. A coil mesh generator script and a field computation script may be further combined into one script and augmented with a parametric loop to enable coil analysis and design (Makarov et al., 2019).

### A5. Software compatibility

The toolkit has been tested on both Windows and Linux platforms using MATLAB 2018a. Generally, no MATLAB toolboxes are necessary with one minor exception. The script `model02_main_script.m` from subfolder `Model` and the function `bemf5_volume_field.m` for computing volumetric fields from subfolder `Engine` both still use a fast built-in function `rangesearch` from the Statistics and Machine Learning Toolbox in order to identify the neighbor facets. This MATLAB toolbox should therefore be installed or a replacement for this function needs to be found.

### A6. Known problems

The toolkit will not run with the older versions of MATLAB. Another difficulty might appear when a script cannot find the proper MATLAB path or the previously loaded data. In this case, it is recommended to start the process over, ideally with the first script `bem0_import_model_data.m` of the main folder as described in Section A2.

